# The Caudate Nucleus Undergoes Dramatic and Unique Transcriptional Changes in Human Prodromal Huntington’s Disease Brain

**DOI:** 10.1101/520312

**Authors:** Filisia Agus, Diego Crespo, Richard H. Myers, Adam Labadorf

## Abstract

The mechanisms underlying degeneration of the specific neurons in the striatum of Huntingon’s Disease (HD) brain are currently unknown. The striatum is massively degenerated in late stage HD, making examination of post-mortem brain tissue from symptomatic individuals problematic. Striatal tissue is largely intact in the brains of asymptomatic HD positive (HD+) gene carriers, but these samples are exceedingly rare. In this study, caudate nucleus (CAU) tissue from two asymptomatic HD+ individuals was subjected to high throughput mRNA sequencing (mRNA-Seq) for comparison with similar datasets from symptomatic HD individuals and healthy controls. The overall transcriptional response in HD+ CAU shares much of the same response observed in HD Brodmann Area 9 (BA9) samples, an area that is relatively spared from significant degeneration. A set of differentially expressed (DE) genes predominantly related to the heat shock response are found in common between brain regions, and show much higher induction in HD+ CAU than HD BA9. The most highly perturbed pathways show near complete agreement when comparing diseased tissue with control, and a random forest classifier predicted that the two HD+ CAU samples strongly resemble HD BA9 and not control BA9. Nonetheless, when genes were prioritized by their specificity to HD+ CAU, a large number of pathways spanning many biological processes emerged. Further comparison of HD+ BA9 with HD BA9 identified genes that may be early responders to disease, and have altered expression in symptomatic individuals. This study presents the first and largest examination of asymptomatic brain gene expression to date, and suggests many new avenues of investigation into the mechanisms underlying neurodegeneration in HD.

## 1 Introduction

Huntington’s Disease (HD) is a devastating neurodegenerative disease caused by an expanded trinucleotide CAG repeat in the HTT gene. The striatum, comprising the caudate nucleus (CAU) and putamen, is the primary affected brain region in HD. As many as 90% of neurons are lost in the striatum, which is massively degenerated in the late stages of the disease. Although other brain regions, such as the cerebellum and cerebral cortex show the hallmarks of HTT protein intranuclear inclusions, they are relatively free of neurodegeneration^1,2^. While studying the striatum directly in post mortem HD brains is preferable, the lack of neurons in these highly degenerated tissues makes interpretation difficult. CAU samples from post-mortem human brains of asymptomatic HD gene positive (HD+) individuals, who died before evidence of significant degeneration has occurred, avoid this difficulty but are extremely rare.

Previously, we performed unbiased transcriptomic analysis with high throughput sequencing (mRNA-Seq) in pre-frontal cortex Brodmannn area 9 (BA9) of twenty HD and forty-nine non-neurological control brain samples^3^. Neuroinflammation and developmental pathways were implicated by the differentially expressed (DE) genes from this study, and there was evidence that every major resident brain cell type (i.e. both neurons and glia) is implicated in HD pathogenesis. However, since all of these individuals were symptomatic and at an advanced stage of disease at the time of death, it was unclear which aspects of the gene expression signature were causes and which were consequences of disease. Examining gene expression from brain tissue of asymptomatic HD+ individuals provides an opportunity to address this key question, as gene expression changes that are present prior to evidence of symptoms and neurodegeneration offer an opportunity to gain insight into initiating disease processes. Furthermore, comparing gene expression changes in BA9 and CAU of the same individuals affords an opportunity to examine how the changes in a relatively unaffected tissue (BA9) reflect those observed in the primarily affected brain region (CAU).

The Myers lab has obtained from the McLean Brain Tissue Resource Center (BTRC), brain tissue from BA9 of three asymptomatic HD+ individuals, as well as CAU from two of these same individuals. These tissues and age and sex matched controls were subjected to mRNA sequencing to assess genome wide alterations in gene expression. The HD+ expression dataset was then compared with our previous HD mRNA-Seq datasets^4^, as well as BA9 and CAU mRNA-Seq samples from the Genotype-Tissue Expression (GTEx) database. The goals of this study were to 1) identify DE genes in the CAU prior to clinical onset and neurodegeneration, 2) compare DE genes between BA9 and CAU in HD+ individuals to identify region-specific and common expression patterns, and 3) identify genes involved in the early vs late disease process.

## 2 Results

Table 1 contains a summary of the datasets used in this study.

**Table 1.** Sample sizes for each class. PMI and Death columns are means followed by standard deviation. Complete sample information is included in Supplemental Table X. HD+ = asymptomatic HD gene positive, HD = symptomatic HD, C = non-neurological control, GTEx = the Genotype-Tissue Expression database, BA9=Brodmann area 9, CAU=Caudate nucleus.

| Sample type | Num samples | PMI | RIN | Age of death |
| --- | --- | --- | --- | --- |
| HD BA9 | 26 | 16.04 ± 7.65 | 7.29 ± 0.89 | 59.77 ± 10.42 |
| HD+ BA9 | 3 | 24.17 ± 8.63 | 7.9 ± 0.62 | 51.33 ± 33.56 |
| C BA9 | 52 | 15.23 ± 9.44 | 7.94 ± 0.64 | 67.88 ± 16.97 |
| GTEx BA9 | 90 | 14.13 ± 4.16 | 7.25 ± 0.87 | 58.63 ± 8.8 |
| HD+ CAU | 2 | 27.97 ± 7.91 | 7.35 ± 0.49 | 67.5 ± 26.16 |
| C CAU | 2 | 31.24 ± 9.64 | 7.8 ± 1.27 | 66.5 ± 21.92 |
| GTEx CAU | 102 | 14.03 ± 4.13 | 7.66 ± 0.76 | 60.07 ± 6.71 |

Five different pair-wise contrasts were performed using the expression estimates, as described in Figure 1 and Table 2. This manuscript will refer to specific analyses by the corresponding numbers in this figure. Analysis (1) compares BA9 for symptomatic HD and neurologically normal controls. Analysis (2) compares HD+ BA9 with C BA9, identifying DE genes likely implicated in the early disease process. Analysis (3) compares HD+ BA9 with HD+ CAU, identifying DE genes caused either by disease or due to differing brain region. Analysis (4) compares HD+ CAU with C CAU, identifying DE genes implicated by the active HD disease process. Analysis (5) compares GTEx BA9 with GTEx CAU, identifying DE genes caused by difference in brain region, to assist in identifying DE genes identified in analysis (3) that are not simply a consequence of different brain region.

**Table 2.** Sample sizes for contrasts performed. First number corresponds to number of samples for column sample type, e.g. for analysis (1) there were 26 HD BA9 and 52 C BA9. The number of DE genes reported have FDR < 0.05. *BA9 control samples that matched the age at death were chosen from the whole control set for this analysis.

| Analysis | Group A | Group B | Sample size | # DE genes | Genes detected |
| --- | --- | --- | --- | --- | --- |
| (1) | HD BA9 | C BA9 | 26 vs 52 | 7789 | 32490 |
| (2) | HD+ BA9 | C BA9 | 3 vs 9 | 229 | 35843 |
| (3) | HD+ CAU | HD+ BA9 | 2 vs 3 | 1199 | 34084 |
| (4) | HD+ CAU | C CAU | 2 vs 2 | 74 | 35554 |
| (5) | GTE<sub>x</sub> BA9 | GTE<sub>x</sub> CAU | 90 vs 102 | 23778 | 34081 |

**Figure 1.**
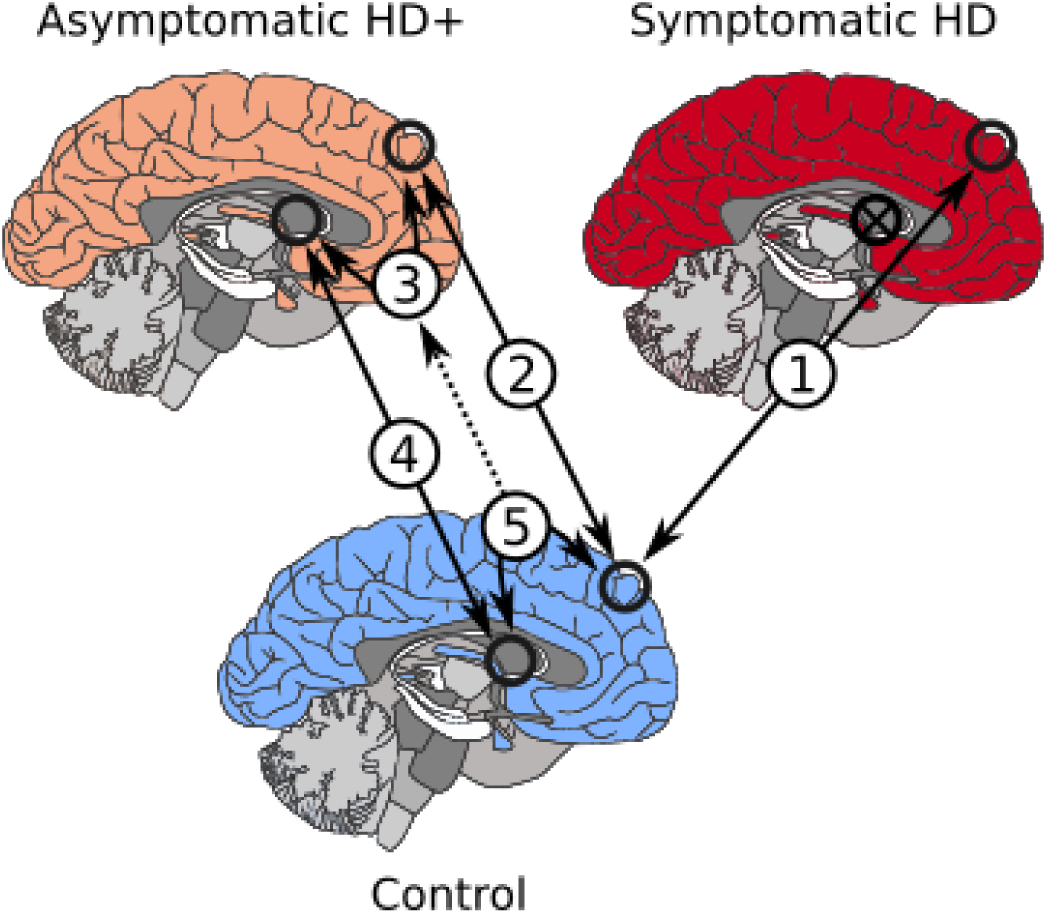
Brain region contrasts performed.

### 2.1 HD BA9, HD+ BA9, and HD+ CAU Show Concordant DE Genes

Figure 2 contains differentially expressed (DE) gene metrics for analyses (1), (2), and (4). In Figure 2A, we see that the fold change distribution is similar between all three analyses, where more genes have increased expression overall than decreased and that this is particularly evident in the HD+ versus control analyses. The overlap of DE genes at FDR < 0.05 in Figure 2B shows that analyses (1) and (4) are more similar to each other than to (2). Figure 2C depicts the similarity in log 2 fold change (L2FC) for the HD+ versus C in BA9 with HD versus C in BA9 (top figure) and the HD+ versus C in CAU with HD versus C in BA9 (bottom figure) for DE genes at *p* < 0.05 in both groups. These two figures show the extent of similarity of L2FC across these different contrasts. It is interesting to note that the symptomatic HD BA9 expression profile is well correlated with the HD+ versus C in CAU (Spearman *ρ* = 0.55) and consequently the HD BA9 appears to be a good model for early disease effects in HD. This concordance is particularly remarkable when considering that the numbers of samples in the HD+ vs C analyses are extremely small.

**Figure 2.**
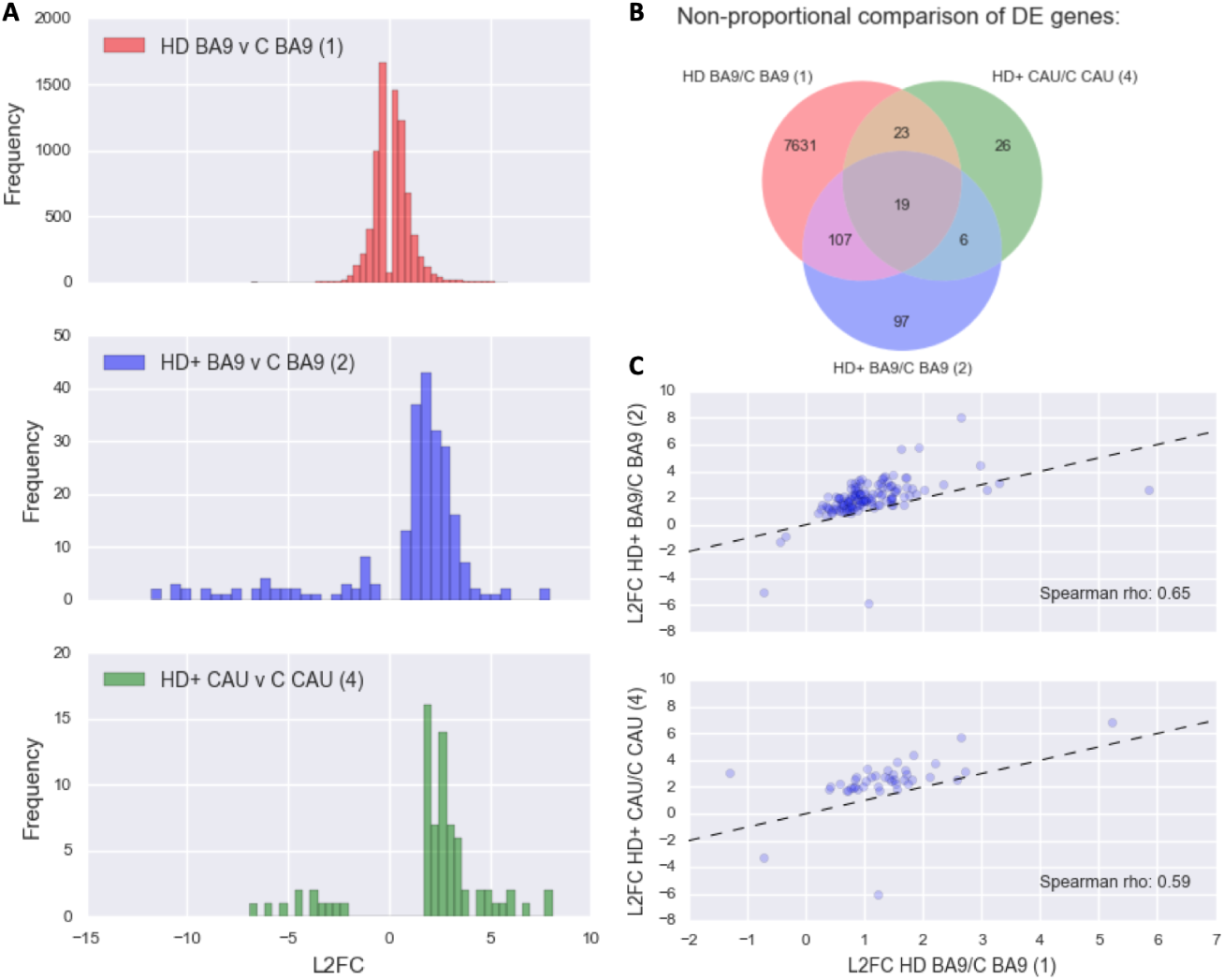
DE Genes from (1), (2), and (4). A) Histograms of log2 fold changes B) 3-way Venn diagram of significant DE genes C) Scatter plots of log2 fold change in DE genes in (1) vs (2) and (1) vs (4) with Spearman *ρ*

The overlapping DE genes in Figure 2B provide insight into both common gene signatures across brain regions and disease state as well as those unique to individual conditions. Table 3 contains the DE statistics for the 19 genes found in the intersection of analyses (1), (2), and (4). These genes are perturbed across the entire disease course, from the HD+ BA9, which is the least affected tissue, to the most severely degenerated HD BA9 samples. All of these genes implicate the neuroinflammatory and neuroimmune responses, and seven of the 19 genes (BAG3, HSPA6, HSPB1, SERPINH1, DNAJB1, HSPA1A, HSPA1B) have direct roles in the heat shock response. As expected, the genes from Table 3 are highly enriched for unfolded protein binding, molecular chaperones and focal adhesion, heat shock response, apoptosis, and response to oxidative stress by DAVID functional enrichment clustering (see Supplementary Table **??**).

**Table 3.**
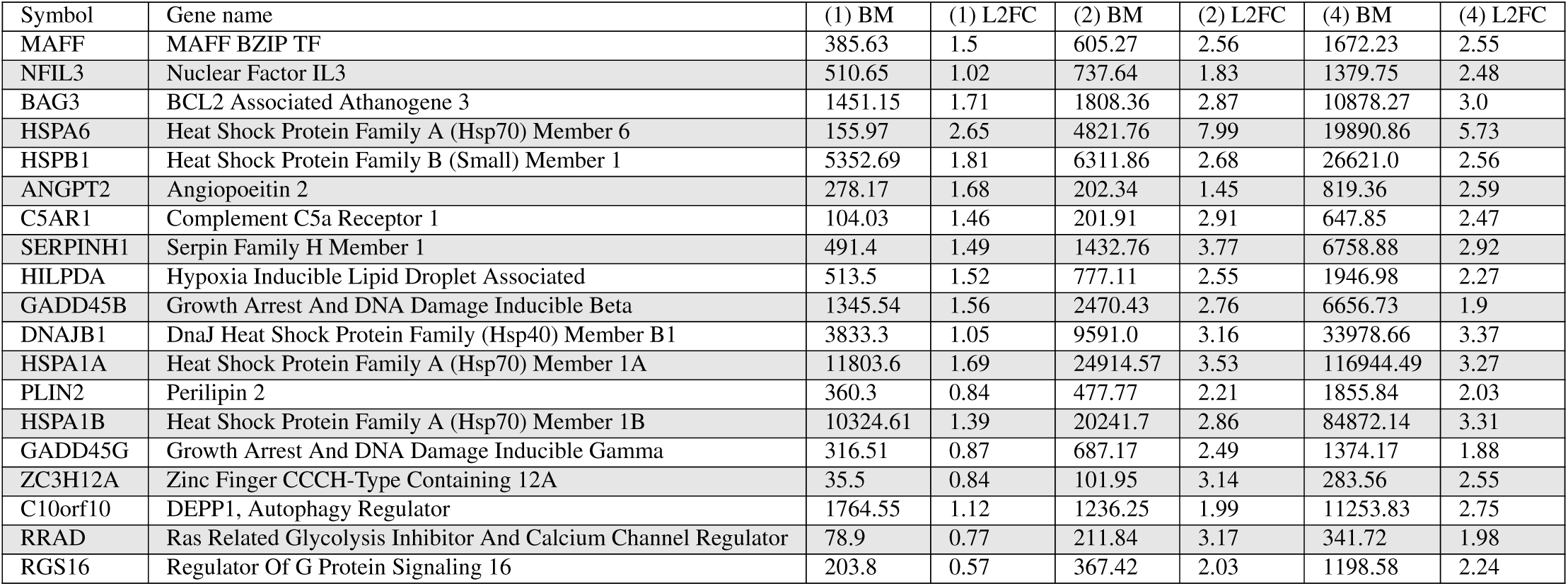
Common response genes in HD BA9, HD+ BA9, and HD+ CAU, corresponds to middle intersection of Venn diagram in figure 2B. Base mean columns are the mean normalized counts from the corresponding analysis. L2FC is log 2 fold change estimated by DESeq2. BM - base mean (number of normalized counts) for the gene.

Figure 3 contains normalized counts distributions for each sample group for the 19 common DE genes from Figure 2B. From left to right in each plot are counts from GTEx BA9, GTEx CAU, C BA9, C CAU, HD BA9, HD+ BA9, and HD+ CAU sample groups. Since there are so few C CAU samples in this study (i.e. only 2), we include the GTEx CAU counts (102 samples) to illustrate that our C CAU counts are well within the expected range for these genes. In every case except ANGPT2, the mean expression level increases from HD BA9 to HD+ BA9 to HD+ CAU. This increase in expression is particularly large for HSPA6, which shows a 256 fold abundance increase in HD+ CAU vs C CAU. Since the HD+ BA9 samples are the least affected tissues of the three disease groups, it is interesting to note that the asymptomatic HD+ BA9 samples show higher expression than the symptomatic HD BA9 samples overall for these genes.

**Figure 3.**
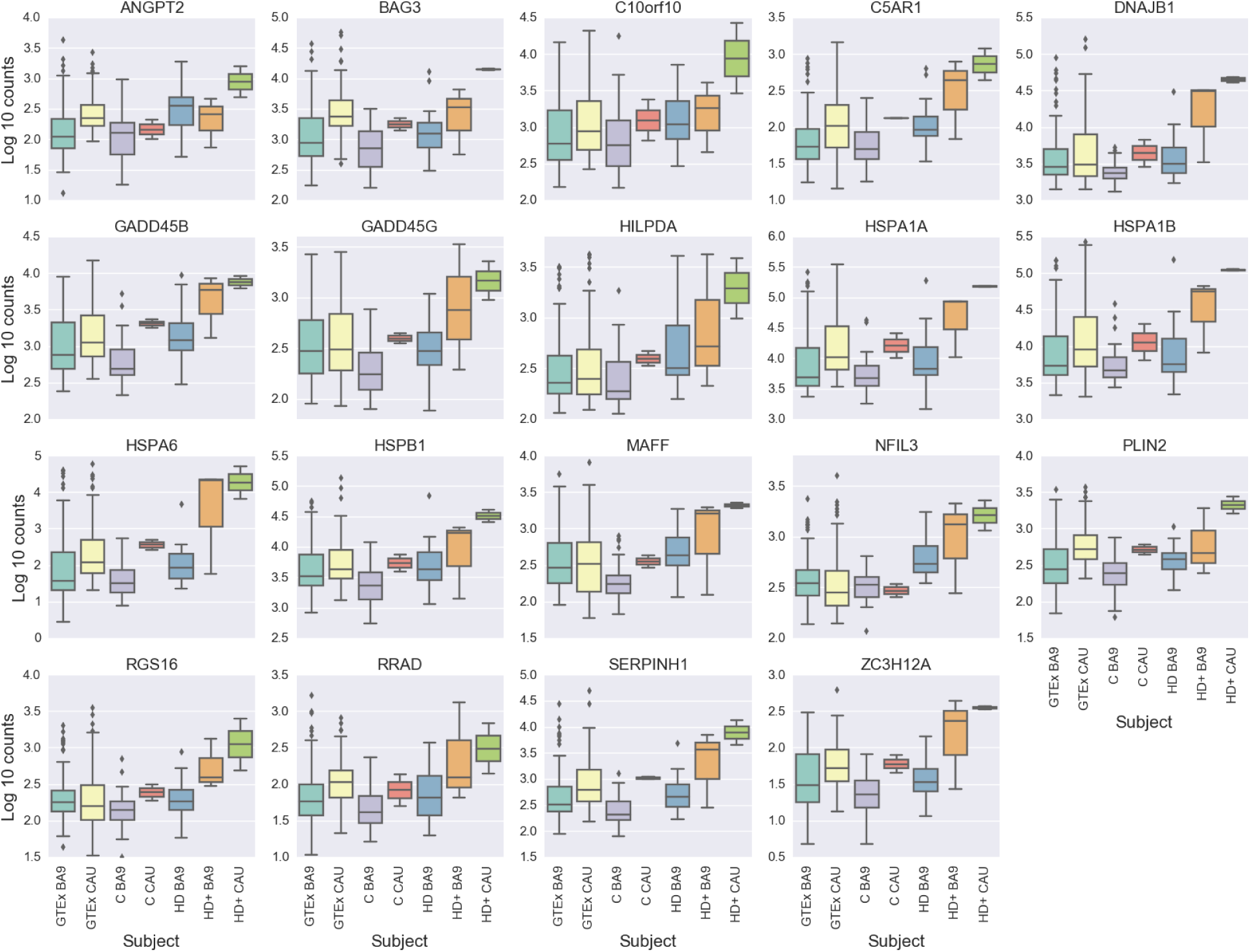
Boxplot of 19 common response genes in all analyses, corresponds to middle intersection of Venn diagram in figure 2B. Base mean columns are the mean normalized counts from all samples. The first two boxes correspond to GTEx BA9 and CAU, respectively, followed by the C BA9 and C CAU from this study. The last three in each plot depict HD BA9, HD+ BA9, and HD+ CAU, respectively.

The 26 genes that are uniquely DE in (4) from Figure 2B (in green segment) appear in Table 4. These genes show only weak functional enrichment for extracellular space compartment, and plasma membrane by DAVID functional enrichment analysis (see Supplementary Table **??**), but we make two remarkable observations. First, several genes are consistent with the heat shock and inflammatory response observed in the common DE genes and in (1) more broadly, including HSPH1, CCL19, and CX3CR1. Second, four of the genes are readthrough transcripts (RPS10-NUDT3, UBE2F-SCLY, RPL17-C18orf32, and RP5-850E9.3) that originate from different chromosomes.

**Table 4.**
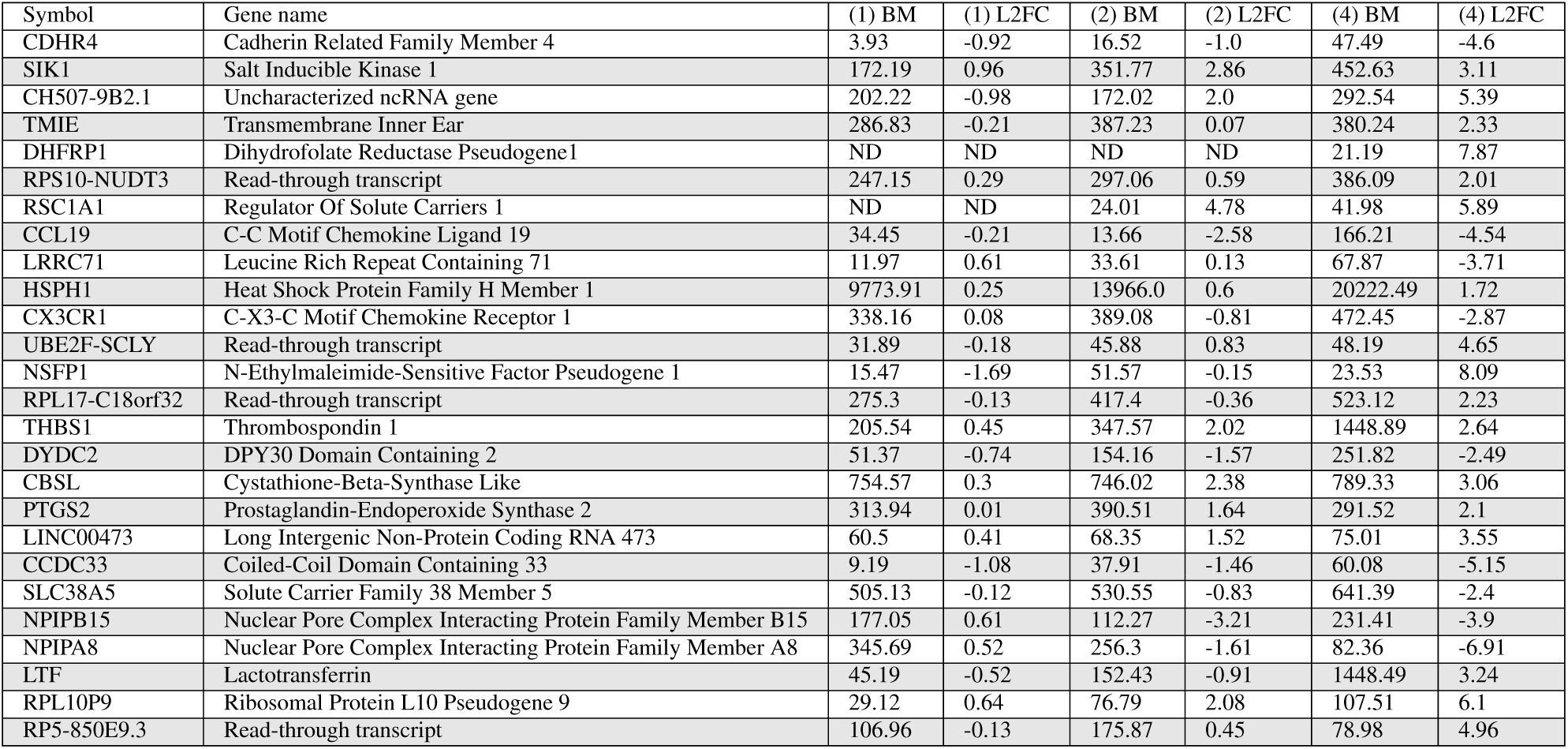
Unique response genes in HD+ CAU, corresponds to only the green area diagram in figure 2B. Base mean columns are the mean normalized counts from the corresponding analysis. L2FC is log 2 fold change estimated by DESeq2. FDR < 0.05 are considered significant for analyses (1) and (2). BM - base mean (number of normalized counts) for the gene. ND - genes not detected or too lowly abundant for consideration in the corresponding samples.

### 2.2 HD+ vs C CAU Enriched Pathways Are A Subset of Those In HD BA9

We next performed gene set enrichment analysis on each DE gene list to identify associated biological functions. Figure 4 contains the result of gene set enrichment analysis from analyses (1), (2), and (4) using the GSEA^5^ algorithm as implemented in the fgsea R package^6^ against the MSigDB C2 Canonical Pathway gene set database^5,7^. Analysis (1), which has the most power to detect DE genes identifies 195 significantly enriched pathways at FDR < 0.05. All 13 of the pathways identified in the HD+ CAU versus C CAU are among these 195 of (1), and eleven of these are also seen in the HD+ BA9 versus C BA9. The substantial overlap of the enriched pathways suggests that the most highly perturbed pathways in the prodromal phase of disease expression are also detected in late stage HD BA9. Only seven pathways, seen in the HD+ BA9 versus C BA9 are not also seen in (1). Table 5 lists the 16 gene sets that are significantly enriched in either both (1) and (4) (9 gene sets) or are unique to (2) (7 gene sets). Consistent with our previous work^4^, the enriched pathways heavily implicate an increase in neuroimmune and neuroinflammatory response, an increase in transcriptional activity, and a decrease in neuron-related pathways.

**Table 5.**
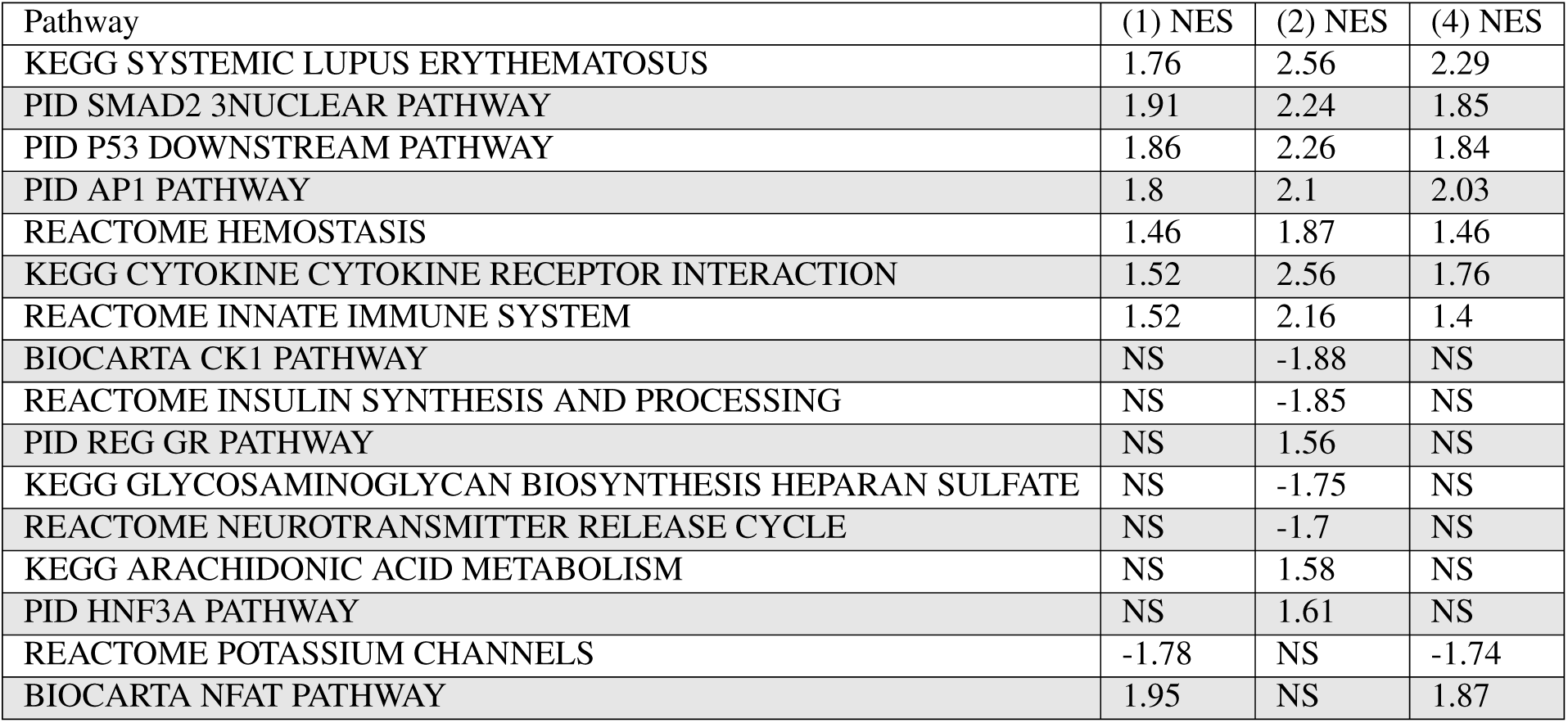
Significantly enriched pathways in intersection of (1) and (4) or unique to (2) from Figure 4. NES = normalized enrichment score from GSEA, where positive or negative values indicate the genes in the pathway are increased or decreased, respectively, in disease compared with control. NS = not significant.

**Figure 4.**
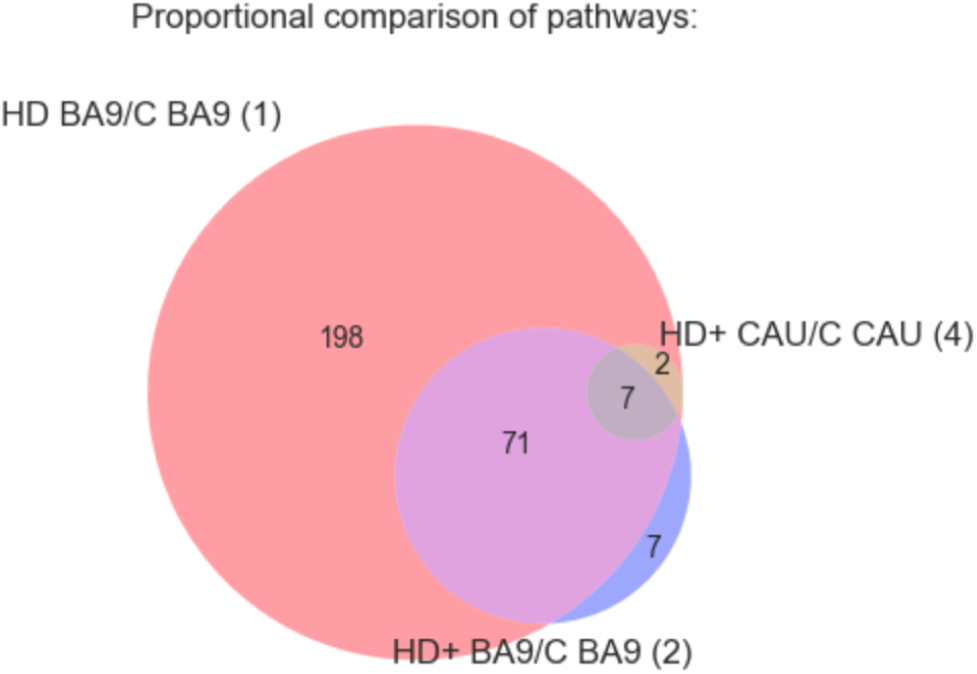
GSEA results for DE genes from analyses (1), (2), and (4). Figure shows overlap of significantly enriched MSigDB C2 Canonical Pathway gene sets at *p*_*ad j*_ < 0.05 irrespective of direction of effect. Selected gene sets are included in Table 5 and the full results are in Supplemental Table 2.

### 2.3 HD BA9 DE Genes Perfectly Predict Disease State in HD+ CAU

Figure 5 shows the normalized counts from all HD, HD+, and C samples for the top 200 genes found to be DE in analysis (1) as a heatmap. A distinctive result from our previous HD work^4^ was that a set of homeotic genes, most notably the HOX gene clusters, were selectively increased in HD compared with C. By inspection, HD+ CAU appears to demonstrate similar homeotic gene expression to HD, suggesting that the disease process in asymptomatic caudate does indeed resemble symptomatic cortex in these samples. HD+ BA9 expression in these genes is less pronounced and more closely resembles C samples, further supporting the hypothesis that the effect of disease on BA9 is reduced in HD+ individuals. The results suggest that HD+ CAU is more similar to HD BA9, and HD+ BA9 is more similar to C.

**Figure 5.**
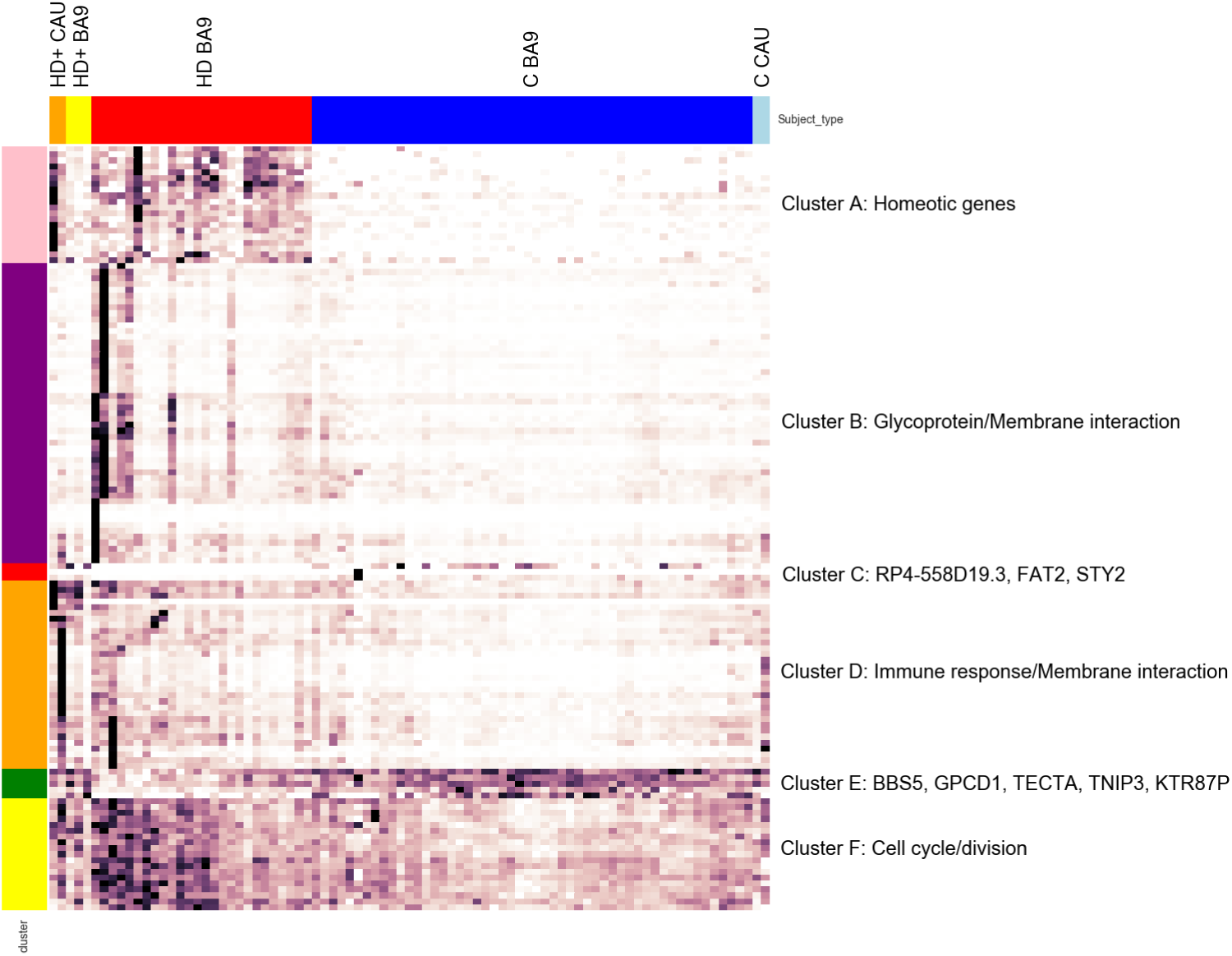
Clustered heatmap of normalized counts of top 200 genes from (1). Row clusters were created manually by inspection guided by clustered dendrogram, and enriched biological pathways (or the genes themselves) for the genes in each cluster are listed as indicated. Color label: HD+ CAU - orange, HD+ BA9 - yellow, HD BA9 - red, C BA9 - blue, C CAU - light blue.

We sought to perform a more unbiased analysis to better quantify the similarity of the HD+ samples to either HD or C by training a random forest decision tree classifier on the HD and C samples. Briefly, a decision tree classifier identifies key features (in our case these are genes) that partition labeled samples (here either HD or C) into like groups using a threshold cutoff for each gene. A decision tree built using a dataset can then be used to predict the class of new samples that were not used to build the tree. To avoid overfitting, the random forest algorithm generates many different decision trees by randomly sampling samples and genes with replacement many times. When applied to a new sample, the output of a random forest decision tree classifier is the number of trees that predicted the sample to have each label. A random forest where all trees classify a new sample to have the same label indicates a perfect classification. A random forest predicting a sample to be of either class with equal frequency has no predictive power. See the Methods section for more details on the random forest decision tree algorithm.

After creating the random forest based on the top 250 significant genes in (1), the forest was used to predict the sample type of each HD+ BA9 and CAU sample. The results of the classifier are in Table 6. Several aspects from the random forest results are of note. First, the random forest perfectly classified both the HD+ CAU samples as symptomatic HD BA9, supporting the intuition built from the heatmap in Figure 5 (Table 6(a)). Second, the HD+ BA9 samples were evenly split between being predicted as HD BA9 and C BA9 (Table 6(a)). This suggests that there are some genes in the HD+ BA9 samples that resemble symptomatic HD BA9, and others that more closely resemble control BA9. We will explore this difference in greater detail in the last section. Third, there is high prediction consistency for HD+ CAU even when choosing 250 genes randomly from (1), and a greater agreement in classifying HD+ BA9 as C BA9 (Table 6(b)). These results suggest that the DE signal for HD+ CAU and in HD BA9 is strong and genome wide, and are consistent with the hypothesis that HD+ BA9 represents a less severe form of the same response as in HD+ CAU and HD BA9. Last, when the model is fully randomized (i.e. random genes and shuffled labels from (1), Table 6(c)), classification consistency is essentially random, consistent with our expectation of the model. Taken together, this unbiased classification analysis supports the hypothesis that changes in BA9 after symptoms have appeared are reflected in the asymptomatic HD+ caudate and, to a lesser degree, in HD+ BA9.

**Table 6.**
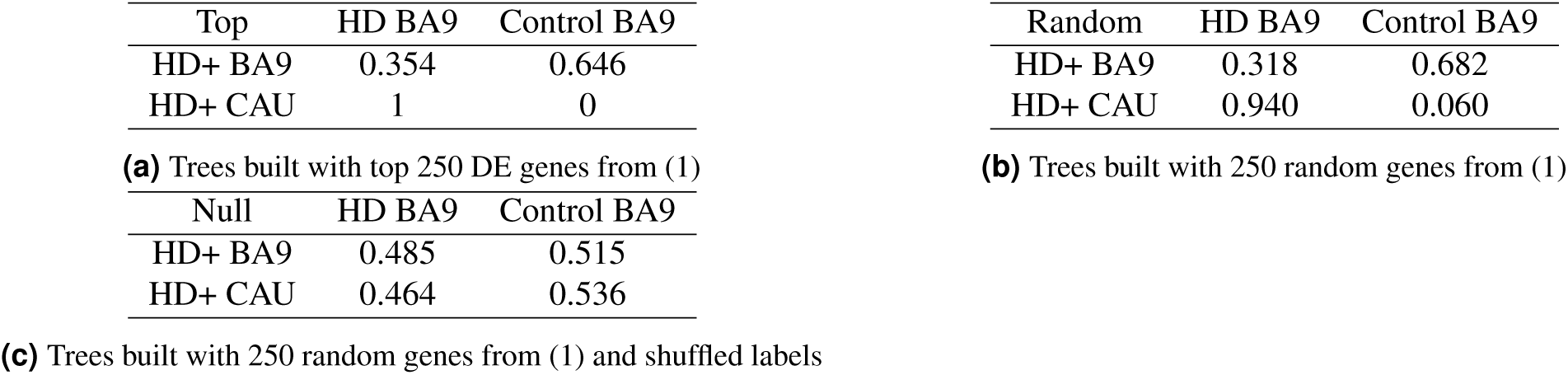
Random forest decision tree classifications of HD+ using genes from (1). All figures are the fraction of 20,000 trees that predicted each sample to have the corresponding label indicated in the column. E.g. 49.5% of the trees predicted HD+ BA9 samples to be HD BA9.

### 2.4 Gene Expression Patterns and Pathways Unique To HD+ CAU

Understanding the factors that cause the caudate to degenerate first in HD is critical in understanding the HD disease process. Due to the small number of HD+ and C CAU samples (2 and 3, respectively), the DE statistics for this direct comparison is likely to be highly influenced by noise, as evidenced by the small number of enriched gene sets in this comparison seen in Figure 4. In addition, directly comparing HD+ CAU and HD+ BA9 may reveal differences between brain regions that are not prominent when comparing the results of corresponding pairwise comparisons. We therefore devised a statistical strategy to identify genes and pathways that are as robust and specific to HD+ CAU as possible.

Since disease status and brain region are convolved in the DE genes identified in (3), we sought to identify genes that differ between CAU and BA9 due to the disease process, and not due to differences in brain region. To accomplish this, the DE results from (3) and (5) were compared by computing a t-statistic of the difference in log2 fold change estimates and their standard errors reported by DESeq2 (see Methods section). In essence, this statistical procedure quantifies the difference in log2 fold change of genes when comparing HD+ CAU versus HD+ BA9 while de-emphasizing genes that are different due to differences in brain region. The resulting statistics allow genes to be ranked by the degree of relevance to the disease process in HD+ CAU. Table 7 contains the top 10 genes ranked by descending absolute value of the t-statistic to illustrate this strategy. For example, CFAP157 is increased 19.6 (2^4.3^) fold in GTEx CAU over GTEx BA9, but is decreased by 1.09 (2^*-*0.13^) fold in HD+ CAU over HD+ BA9, resulting in an difference in fold change of −4.43 (i.e. 0.13 4.3 = 4.43). TVP23C-CDRT4, another readthrough transcript, is essentially unchanged in GTEx CAU compared with GTEx BA9, but is increased 5.85 fold in HD+ CAU over HD+ BA9 (2.55 *-* (*-*0.02) = 2.57).

**Table 7.** Top 10 genes that show different effect sizes (L2FC) between (3) and (5). These genes are most likely perturbed in CAU specifically due to HD and not due to brain region. Δ L2FC is the log2 fold change of (5) minus (3), where a positive value means that gene expression is greater in HD+ CAU vs BA9 than GTEx CAU vs BA9. Full results are in Supplemental Table 4.

| ENSGID | Gene Symbol | HD+ L2FC (3) | GTE<sub>x</sub> L2FC (5) | $\Delta$ (3) vs (5) L2FC | *t* |
| --- | --- | --- | --- | --- | --- |
| ENSG00000160401 | CFAP157 | -0.13 | 4.30 | -4.43 | 20.45 |
| ENSG00000077327 | SPAG6 | -2.20 | 2.64 | -4.84 | 15.28 |
| ENSG00000152611 | CAPSL | -2.59 | 3.65 | -6.24 | 14.63 |
| ENSG00000181085 | MAPK15 | -1.65 | 2.71 | -4.36 | 12.89 |
| ENSG00000154914 | USP43 | -3.32 | -0.80 | -2.52 | 12.87 |
| ENSG00000169436 | COL22A1 | -3.82 | -0.31 | -3.51 | 12.82 |
| ENSG00000259024 | TVP23C-CDRT4 | 2.55 | -0.02 | 2.57 | -12.75 |
| ENSG00000162747 | FCGR3B | 3.21 | 0.31 | 2.90 | -12.45 |
| ENSG00000118113 | MMP8 | 4.70 | 0.13 | 4.57 | -12.43 |
| ENSG00000103569 | AQP9 | 1.59 | -0.72 | 2.31 | -12.41 |
| ENSG00000140795 | MYLK3 | -0.78 | 1.87 | -2.65 | 12.10 |

The resulting t-statistics from this analysis induced a ranking of genes that were then subjected to gene set enrichment analysis against the MsigDB C2 Canonical Pathway gene set database. The analysis identified 405 significantly enriched gene sets at FDR < 0.05, and all but one of these gene sets were positively enriched, indicating that genes increased in HD+ CAU relative to HD+ BA9 have strong functional coherence (full fgsea results in Supplemental Table 3). These results were combined with the enriched gene sets from (1), (2), and (4) and subsequently divided into so-called Agreement Classes based on the pattern of significance across all four analyses. The Agreement Class is an ordinal indicator for the degree of HD+ CAU-specificity as follows. CAU Unique gene sets are only seen in HD+ CAU relative to HD+ BA9 (i.e. (3) vs (5)). CAU Enhanced are enriched gene sets in HD BA9 vs C BA9 (1) as well as in either HD+ CAU vs C CAU (4) or HD+ CAU relative to HD+ BA9 ((3) vs (5)). Finally, BA9 Unique only show enrichment in HD BA9 vs C BA9 (1). To aid in interpretation, the gene sets were manually curated into 10 high level functional categories: Angiogenesis/Blood Brain Barrier (BBB), Apoptosis, Cell Cycle/Development, Cytoskeleton/Extracellular Matrix (ECM), Immune Response/Cancer, Metabolism, Neuron System, Protein Folding/Other, Signaling, and Transcription/Translation. To illustrate these ideas, a heatmap of the enriched gene sets related to the Neuron System is in Figure 6A.

**Figure 6.**
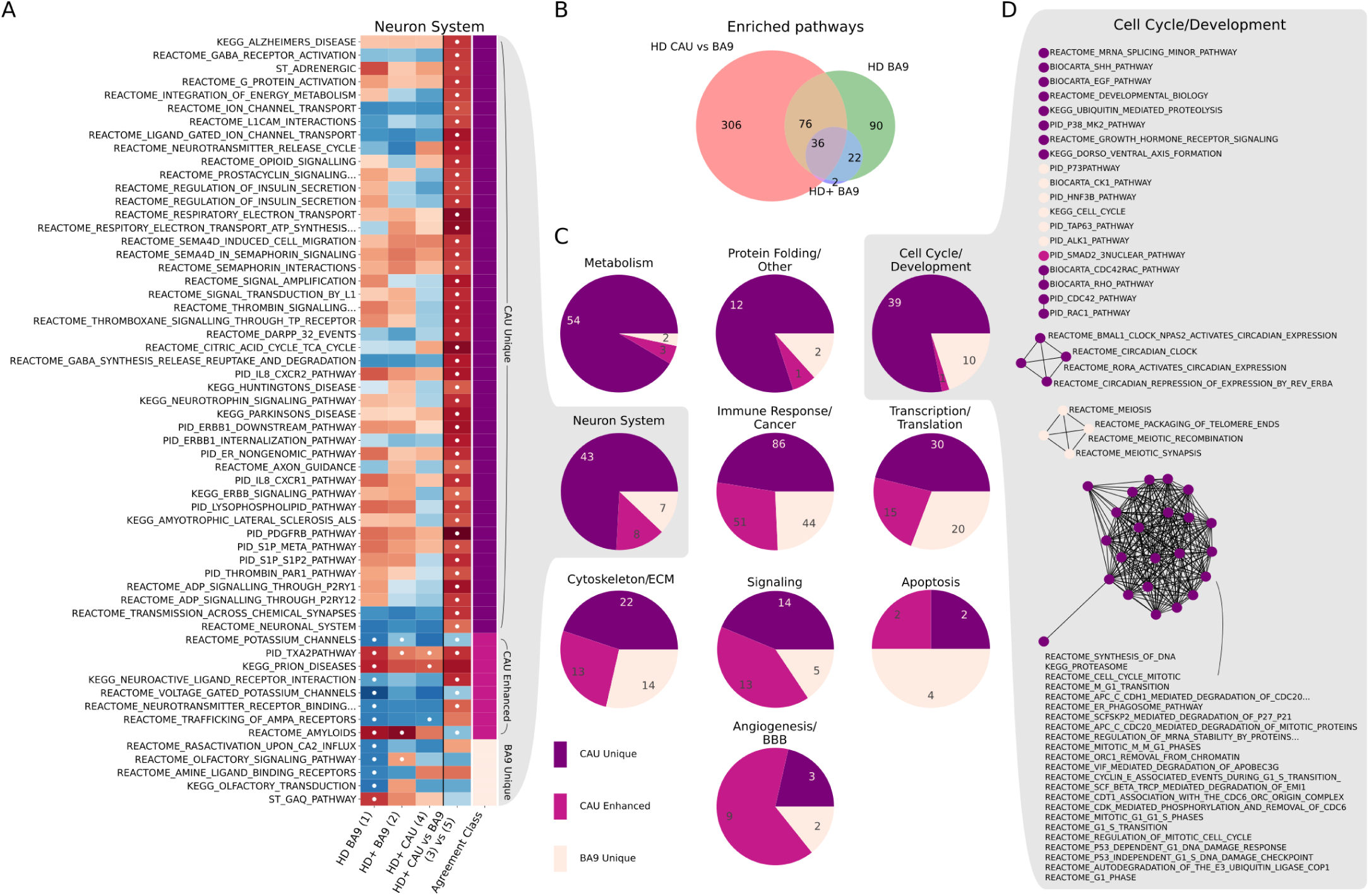
A) Enriched gene sets related to the Neuron System. The first four columns plot Normalized Enrichment Score (NES) of GSEA analyses from (1), (2), (4), and (3) vs (5), respectively, where red and blue correspond to positive and negative NES scores, respectively. The fifth column indicates the Agreement Class of each gene set, assigned according to HD+ CAU-specificity. Cells with white dots indicate that gene set is significantly enriched in the corresponding analysis. B) Overlap of significantly enriched gene sets regardless of category. The gene sets enriched in (4) are a subset of those in (1), and thus are not listed. C) Distribution of gene sets by agreement class divided into ten high level functional categories, showing that some functions are more selectively enriched in HD+ CAU relative to BA9 than others. D) Graph-based representation of the Cell Cycle/Development gene sets from C. Each node is a gene set, and nodes with connected edges share more than 25% of their leading edge genes, thus representing the same expression signal.

As seen in Figure 6B, 306 out of 405 significantly enriched gene sets are unique to CAU relative to BA9. The distribution of these unique gene sets varies by biological process (Figure 6C), where processes related to Cell Cycle/Development, Metabolism, Neuron System, and Protein Folding/Other show the greatest proportion of CAU-unique gene sets. This is in contrast to Angiogenesis/BBB, where most of the gene sets are seen in both CAU relative to BA9, and in BA9 independently. Gene sets related to Apoptosis, Cytoskeleton/ECM, Immune Response/Cancer, Signaling, and Transcription/Translation have a mixture of CAU unique, CAU enhanced, and BA9 specific gene sets. When we examine the enriched gene sets from Cell Cycle/Development more closely using a graph-based representation (Figure 6D), we observe that there are two distinct groups of genes enriched separately in CAU vs BA9 and BA9 itself. In particular, BA9 is enriched for a set of genes relating to meiosis, whereas the CAU unique processes involve mitosis. Heatmaps and graph representations of all other categories are included in Supplemental File **??**. Overall, this comparison of enriched gene sets in HD+ CAU relative to BA9 with the other two brain regions identifies the common and different cellular processes that are active in different brain regions.

### 2.5 Comparing DE Gene Lists Identifies Early vs Late Responding Genes

In the random forest analysis discussed above, we noted that the HD+ BA9 samples were classified either as HD BA9 or C BA9 with approximately equal frequency. This suggests that there are some genes with an expression pattern that resembles HD BA9 and some that are yet unaffected in asymptomatic HD+ BA9. Thus, the genes that are consistent between HD+ BA9 and HD BA9 are genes that may form an early response in HD, whereas the genes whose expression differs from HD BA9 might still be intact and only respond later in the disease. We sought to identify which genes were early vs late responders by applying our t-statistic strategy comparing log2 fold changes between analyses (1) and (2). Table 8 contains results from the t-statistic based analysis of (1) and (2).

**Table 8.** Genes partitioned by significance within analyses (1) and (2) and fold change difference between these analyses. Numbers in parentheses are genes that appeared in the corresponding analysis but were filtered out in the other.

|  | (1) HD vs C BA9 |  | (2) HD+ vs C BA9 |  |  |
| --- | --- | --- | --- | --- | --- |
|  | DE | Not DE | DE | Not DE | Total |
| Sig. Between | 4454 | 11670 | 218 | 15906 | 32248 |
| Not Sig. Between | 3281 (54) | 12634 (387) | 9 (2) | 15906 (3765) | 31830 |
| Total | 7735 | 24304 | 227 | 31812 |  |
| Grand total | 32039 |  | 32039 |  |  |

Of particular note are the 215 genes that are DE in (2) and have fold changes different from (1). These are the genes that may reflect early disease processes not identifiable in symptomatic individuals post mortem. We extracted these 215 genes and plotted their log fold changes to examine the relationship between groups as depicted in Figure 7. Genes in quadrants I and III of the figure are genes that show the same direction of effect (i.e. up or down) but have a different size of effect. Genes in quadrants II and IV are genes that show differential behavior, and are thus potentially unique responses early in the disease process that are not observed in symptomatic HD BA9.

**Figure 7.**
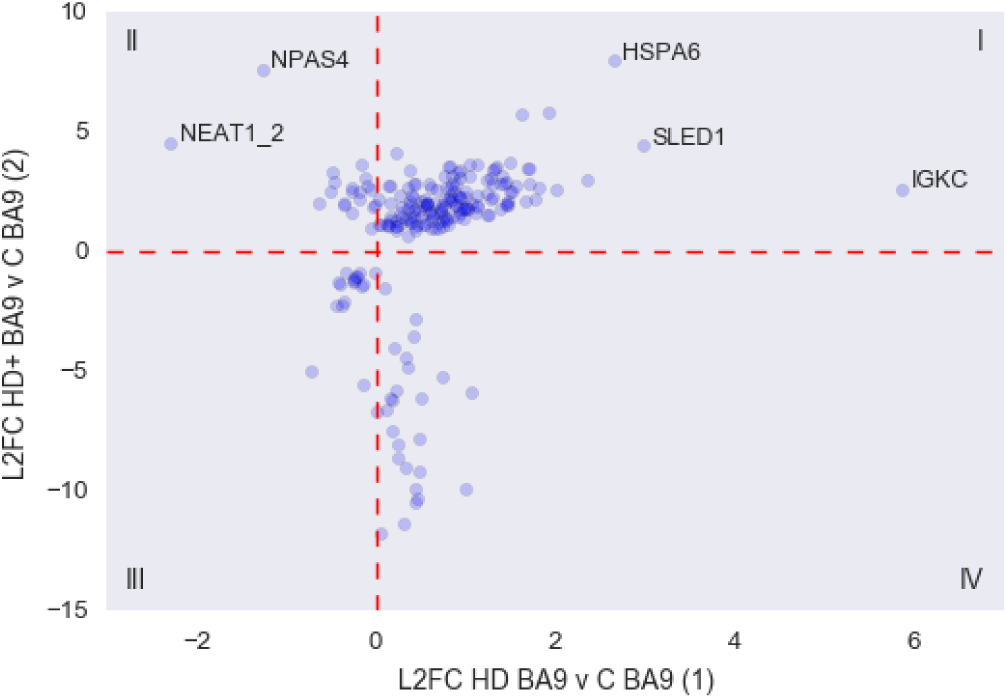
Scatter plot of fold changes from 215 early response genes from (1) and (2). Still need to label names of the genes in II and IV

The statistics from the genes in quadrants II and IV, as well as the 4 additional genes detected in (2) that were filtered out of (1) due to low counts are listed in Tables 9 and 10. Most of the genes that are down regulated in (2) with respect to (1) are ribosomal protein genes that are essentially absent from HD+ BA9. The two genes that are massively increased in (2) but decreased in (1) are NPAS4, Neuronal PAS Domain Protein 4, and NEAT1 2.

**Table 9.** Putative early response genes in HD+ BA9 from figure 5 quadrant II. Base mean columns are the mean normalized counts from the corresponding analysis. L2FC is log 2 fold change estimated by DESeq2.

| Ensembl ID | Gene name | (1) Basemean | (1) L2FC | (2) Basemean | (2) L2FC |
| --- | --- | --- | --- | --- | --- |
| ENSG00000125740 | FOSB | 363.46 | -0.17 | 1438.94 | 3.63 |
| ENSG00000174576 | NPAS4 | 165.91 | -1.27 | 2473.17 | 7.64 |
| ENSG00000278050 | NEAT1_2 | 2.51 | -2.29 | 14.4 | 4.57 |
| ENSG00000135625 | EGR4 | 263.89 | -0.48 | 832.95 | 3.35 |
| ENSG00000173391 | OLR1 | 229.69 | -0.05 | 566.64 | 2.62 |
| ENSG00000198576 | ARC | 897.43 | -0.51 | 2278.26 | 2.54 |
| ENSG00000158050 | DUSP2 | 308.83 | -0.28 | 850.53 | 2.54 |
| ENSG00000153234 | NR4A2 | 519.76 | -0.18 | 1049.01 | 2.21 |
| ENSG00000120738 | EGR1 | 1826.3 | -0.64 | 3887.13 | 2.06 |
| ENSG00000248713 | RP11-766F14.2 | 31.18 | -0.37 | 71.78 | 1.96 |
| ENSG00000160223 | ICOSLG | 796.19 | -0.36 | 813.93 | 2.03 |
| ENSG00000232352 | SEMA3B-AS1 | 30.91 | -0.27 | 33.01 | 1.67 |
| ENSG00000174429 | ABRA | 14.35 | -0.11 | 35.74 | 2.78 |
| ENSG00000273186 | RP11-339B21.10 | 9.21 | -0.46 | 10.63 | 2.95 |
| ENSG00000122877 | EGR2 | 136.81 | -0.27 | 366.35 | 2.72 |
| ENSG00000123358 | NR4A1 | 1771.51 | -0.19 | 4397.13 | 2.52 |
| ENSG00000162783 | IER5 | 1079.98 | -0.02 | 2118.33 | 2.0 |
| ENSG00000105722 | ERF | 968.7 | -0.05 | 1248.28 | 1.01 |
| ENSG00000244062 | RP11-404G16.2 | 31.01 | -0.13 | 52.64 | 3.11 |
| ENSG00000184378 | ACTRT3 | 37.62 | -0.05 | 81.25 | 1.9 |

**Table 10.** Putative early response genes in HD+ BA9 from figure 5 quadrant 4. Base mean columns are the mean normalized counts from the corresponding analysis. L2FC is log 2 fold change estimated by DESeq2.

| Ensembl ID | Gene name | (1) Basemean | (1) L2FC | (2) Basemean | (2) L2FC |
| --- | --- | --- | --- | --- | --- |
| ENSG00000258017 | RP11-386G11.10 | 7775.65 | 0.25 | 3679.44 | -8.64 |
| ENSG00000176868 | RP11-334J6.7 | 2651.11 | 0.43 | 1455.75 | -10.47 |
| ENSG00000255082 | GRM5-AS1 | 2209.09 | 0.19 | 1407.72 | -6.19 |
| ENSG00000254873 | RP11-770J1.5 | 980.14 | 0.18 | 620.42 | -7.46 |
| ENSG00000267469 | AC005944.2 | 2028.65 | 0.43 | 1058.48 | -9.89 |
| ENSG00000269604 | AC005523.2 | 1008.72 | 0.06 | 565.69 | -11.78 |
| ENSG00000272379 | RP1-257A7.5 | 697.88 | 0.47 | 385.27 | -7.77 |
| ENSG00000225339 | RP11-513I15.6 | 5979.33 | 0.11 | 3352.43 | -6.56 |
| ENSG00000265401 | RP11-138I1.4 | 4353.79 | 0.23 | 2305.67 | -5.75 |
| ENSG00000232940 | HCG25 | 272.52 | 0.42 | 137.87 | -3.5 |
| ENSG00000271127 | LL22NC03-N64E9.1 | 26.44 | 0.34 | 22.19 | -4.43 |
| ENSG00000273489 | RP11-180C16.1 | 1300.83 | 0.5 | 737.27 | -6.13 |
| ENSG00000228748 | RP13-39P12.3 | 301.31 | 0.44 | 190.55 | -2.79 |
| ENSG00000233427 | RP1-212P9.3 | 28.1 | 0.75 | 13.53 | -5.25 |
| ENSG00000269145 | AC007192.6 | 761.47 | 0.3 | 390.97 | -11.38 |
| ENSG00000279753 | AC011558.5 | 15.79 | 1.07 | 6.54 | -5.88 |
| ENSG00000261641 | LA16c-390E6.5 | 296.63 | 0.99 | 132.13 | -9.85 |
| ENSG00000268220 | RP11-379K17.12 | 505.97 | 0.33 | 317.65 | -8.97 |
| ENSG00000269243 | CTD-2231E14.8 | 214.65 | 0.15 | 109.74 | -6.07 |
| ENSG00000267436 | AC005786.7 | 95.9 | 0.24 | 43.6 | -8.07 |
| ENSG00000279767 | AL513523.2 | 642.42 | 0.46 | 288.95 | -10.33 |
| ENSG00000258430 | RP11-982M15.2 | 210.57 | 0.47 | 103.36 | -9.17 |
| ENSG00000219410 | RP4-761J14.8 | 363.25 | 0.36 | 195.65 | -4.79 |
| ENSG00000120992 | LYPLA1 | 625.77 | 0.1 | 563.93 | -1.49 |
| ENSG00000256341 | RP11-21A7A.3 | 23.56 | 0.21 | 15.33 | -4.01 |
| ENSG00000249141 | RP11-514O12.4 | Na | Na | 4.87 | 5.35 |
| ENSG00000279909 | AC110615.1 | Na | Na | 187.8 | -10.4 |

## 3 Discussion

To the authors knowledge, this is the first genome-wide transcriptome analysis of post-mortem asymptomatic HD+ BA9 and CAU. It is also the first systematic comparison of post-mortem symptomatic (HD) BA9 with asymptomatic (HD+) BA9 and CAU gene expression. Differential expression (DE) analysis identified many genes that show altered abundance between diseased and control tissue across brain regions, and there is a high degree of concordance in the direction of effect for these genes. The genes that are commonly DE in HD BA9, HD+ BA9, and HD+ CAU are strongly enriched for heat shock response, while the DE genes specific to HD+ CAU contain some heat shock elements and read-through transcripts. Gene set enrichment results show a high degree of agreement between these three analyses.

The analysis comparing HD+ CAU to HD+ BA9, and filtered using GTEx data for genes specific to brain region, identified a strikingly large number of significantly enriched gene sets that suggest processes related to Cell Cycle/Development, Metabolism, Neuron System, and Protein Folding are the most uniquely perturbed in the HD+ CAU disease process. Overall this analysis suggests that while a large proportion of disease processes are shared between CAU and BA9, there are distinct and important sets of genes perturbed in each brain region related to the disease process. Nonetheless, when the HD+ samples are classified using a random forest classifier built using the symptomatic BA9 samples, there is complete consensus that HD+ CAU most closely resembles HD BA9, while HD+ BA9 resembles aspects of both diseased and control brain. The homeotic and inflammatory gene signatures appear to be equally present in the HD+ CAU and HD BA9, suggesting a similar process affects the cellular milieu in both brain regions.

Finally, we identified key genes that appear to be early responders to the disease process by comparing HD+ BA9 and HD BA9. HD+ BA9 appears to be the least affected brain region of the three studied here; therefore, genes that show different behavior in these regions are likely to be part of an early response that is lost as the disease process progresses. The two genes in particular that are increased in HD+ BA9 relative to HD BA9 are NPAS4, which as been implicated in the cortex of mouse models of HD^8^ and NEAT1, which has been shown to be associated with neuronal hyperactive state^9^. A second group of poorly annotated but consistently expressed genes seems to be uniquely expressed in HD BA9 and may be evidence of severe transcriptional dysregulation previously observed in this tissue^3,4^.

Despite the small HD+ sample size, the consistency between the HD+ and HD BA9 results supports the robustness of these findings. Not only do the overall effect size and enriched pathway signatures agree to a great extent, many of the biological processes implicated are well supported in the literature. Immune response has been heavily implicated in HD and neurodegenerative disease in general^3,4,10–14^, and the broad agreement between the diseased tissues across brain regions in this study lends support to the role of inflammation in the prodromal HD brain. Of particular note is the common heat shock response observed in the common DE genes in all comparisons with control. The heat shock system is primarily responsible for maintaining proteostasis and protein conformation during times of stress, and has been directly implicated in both animal^15^ and in vitro^16^ models of Huntington’s disease. The fact that expression of key heat shock genes appears to be perturbed across the entire disease course is strong evidence of the important role these proteins play in disease.

The differences revealed between HD+ CAU and HD+ BA9 may offer insight into why the striatum is uniquely vulnerable to neurodegeneration. The enriched functional categories that are the most specific to HD+ CAU include Metabolism and Cell Cycle and Development. Interestingly, when the cell cycle gene sets are examined closely (Figure 6D), we observe that the gene sets uniquely enriched in HD+ CAU are related to mitosis, while the smaller number enriched in BA9 involve meiosis. The striatum, unlike the cortex, has a resident population of neuroblasts that enables neurogenesis in the adult human brain^17^. A recent hypothesis has proposed that these neuroblasts are impaired in HD, resulting in a lack of repleneshing neurons over time and eventual destruction of tissue^18^. The unique presence of increased mitotic gene expression, paired with the observation that many neuronal pathways are also increased in HD+ CAU compared with HD+ BA9, is strong evidence that neurogenesis is indeed active in this region prior to symptom onset. However, it still remains to be shown why these specific neurons degenerate in the first place, and why this neurogenesis ceases over time. The enrichment of meiosis in BA9 is curious, and does not lend an immediate interpretation. One possibile explanation is that the same signals that trigger neurogenesis in CAU are also present in BA9, but that cortical neurons lack neurogenic capabilities and ‘misfire’ in response to the developmental signals. An intriguing feature of the HD BA9 samples is the expression of homeotic and developmental genes, which might be a consequence of a neuron that is trying to regenerate but cannot.

Given the extreme rarity of HD+ CAU samples, it is difficult to conceive of validation experiments to test these findings given our current disease models. The combined complexity of the central nervous and immune systems makes accurate models of human HD challenging to devise, since there is clear involvement and interaction of major players in both of these systems. Only a few of the many findings in this study have been discussed in this manuscript, and much greater insight may likely be gained from further examination by those specializing in different aspects of the biology implicated here. These results are therefore put forward as a source of hypothesis and inspiration for new models and avenues of research.

## 4 Methods

### 4.1 Human Subjects

The individuals in this study are exempt as defined by the Boston University School of Medicine Institutional Review Board, due to the fact that all analyses were derived from postmortem brain tissue.

### 4.2 Sample processing

26 symptomatic HD and 56 control BA9 mRNA-Seq libraries were used as previously described^4^. Paired BA9 and CAU tissues from two asymptomatic HD gene positive individuals, one additional asymptomatic HD gene positive BA9 sample, and two CAU samples from neurologically normal controls were extracted and processed to generate mRNA-Seq libraries following the procedure previously described^4^. Statistics for new samples reported in this study are found in Table 11. Raw and processed read data have been deposited into GEO under accession GSMXXXXXX.

**Table 11.**
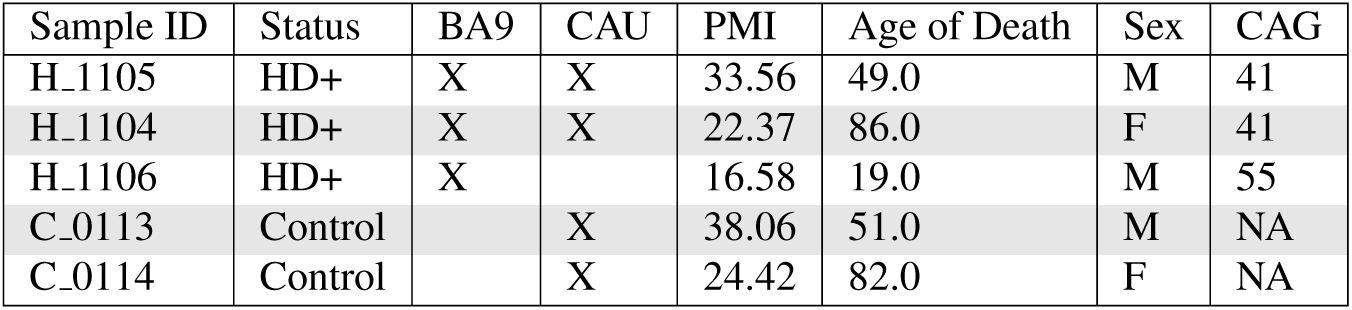
Sample statistics. HD+ are asymptomatic gene positive individuals. Two HD+ individuals had both BA9 and CAU brain tissues available for analysis. Full sample statistics are included in Supplemental Table S1.

| Sample ID | Status | BA9 | CAU | PMI | Age of Death | Sex | CAG |
| --- | --- | --- | --- | --- | --- | --- | --- |
| H_1105 | HD+ | X | X | 33.56 | 49.0 | M | 41 |
| H_1104 | HD+ | X | X | 22.37 | 86.0 | F | 41 |
| H_1106 | HD+ | X |  | 16.58 | 19.0 | M | 55 |
| C_0113 | Control |  | X | 38.06 | 51.0 | M | NA |
| C_0114 | Control |  | X | 24.42 | 82.0 | F | NA |

### 4.3 Quality Control and mRNA Abundance Estimation

mRNA-Seq libraries were subject to quality control and analysis using a custom pipeline. All sequencing libraries were quality- and adapter-trimmed with trimmomatic^19^, and then assessed to be of high quality using fastqc^20^ and MultiQC^21^. Trimmed reads were analyzed with salmon^22^ to obtain mRNA abundance estimates using the GENCODE v26 gene annotation^23^. Abundance estimates from all samples were concatenated into a single matrix and normalized with the DESeq2 normalization method^24^. The normalized expression matrix was investigated for outlier samples using PCA, where no outliers were found (Supplemental File S1 Figure 1). Due to the different numbers of samples, genes in each analysis were filtered using different strategies. For analyses 1, 2 and 5, genes with more than 50% zero counts within each group was filtered out. For analyses 3 and 4 genes with more than 2 zeros and genes with more than 4 zeros was filtered out, respectively. Therefore, the genes detected for each analysis were different as seen in Table 2.

### 4.4 GTEx Analysis of BA9 vs CAU

Post mortem human brain samples from BA9 and CAU brains available from the GTEx project^25^ were downloaded and processed as above. After processing the samples through the quality control pipeline described above, 56 samples were removed due to differences in per base Sequence Content, over representation of sequences, or discrepancies in read length, leaving a total of 90 BA9 samples, 102 CAU samples for analysis. These 192 samples were used to form the basis of a contrast between HD+ BA9 and HD+ CAU samples.

### 4.5 Differential Expression Analysis

Five differential expression contrasts were conducted in this study as described in the analysis matrix of Table 2 and Figure 1. Differential expression statistics for all five analyses were assessed using DESeq2^24^, modeling counts as a function of either disease status or brain region, adjusting for age at death and sex. Differentially expressed genes were considered significant if they had FDR < 0.05.

### 4.6 Gene set enrichment

Gene set enrichment analysis for all DE gene lists was performed using the fgsea^6^ R package in bioconductor^26^ and MSigDB C2 Canonical Pathway database version 6.2^5,7^. GSEA statistics were calculated using each gene list sorted by descending log2 fold change, and significance was assessed for gene sets at FDR < 0.05.

### 4.7 Random Forest Predictive Model to Classify HD+

A random forest of decision trees was used to classify the HD+ CAU and BA9 samples as either HD BA9 or C BA9. A decision tree is a predictive model that iteratively bifurcates a set of labeled samples by identifying features (e.g. genes) that have predictive power when partitioning samples by a fixed threshold. The decision tree algorithm is a machine learning technique that is used to identify features and their levels that best partition a sample set according to given labels. For example, if gene A is expressed between 10 and 20 in one set of samples and between 30 and 40 in another, samples with an expression value less than 25 are likely to belong to one class, while samples with an expression value greater than 25 will belong to the other. If a single gene cannot perfectly divide samples into their labels, additional genes are chosen in a hierarchical fashion until samples with different labels are perfectly partitioned. Once a decision tree has been trained, it may be used to classify previously unobserved samples into the labels used in training.

Individual decision trees trained with all samples are often over-fit, so a randomization technique called random forests are used with decision trees to identify robust predictive features cite?. Random forests perform bootstrap sampling on samples and random selection of features to build a large number of decision trees, where each is a different predictive model with different sets of features. After training, unobserved samples are applied to each decision tree in the forest and the predicted label of each is recorded and reported. The agreement of predicted labels across all trees in the forest is an indication of the predictive power of the overall dataset. A random forest where all trees classify a new sample to have the same label indicates a perfect classification. A random forest predicting a sample to be of either class with equal frequency has no predictive power.

We trained a random forest of decision trees using the HD BA9 vs C BA9 normalized counts matrix to arrive at a predictive model of genes that well classify the samples. The random forest was trained using cross-validation, where the samples are divided into training and test sets. Decision trees in the forest are built using the training samples and their predictive accuracy is assessed on the test set. In this way, cross validation enables assessing the robustness of a classifier and avoids over-fit predictive models. Each random forest contained 20,000 trees, 250 genes, 75% training sets were created with ratios of HD and Control samples which mirrored ratios present in the dataset. Prediction accuracy was assesses as the mean true positive predictions divided by the number of trees across all samples in the test set. 1000 random forests were trained in this way, and the average and standard deviation of true positive rates were recorded for each. See Table 13 for statistics on cross validation prediction accuracy.

**Table 12.** Random Forest Results

**Table 13.** Accuracy Results. Random forest cross validation prediction accuracy statistics for the HD vs C samples. The Random 250 random forests were generated by selecting from a random subset of 250 genes from the overall dataset. The Null random forests were generated by shuffling sample labels and choosing 250 genes at random.

|  | Top 250 DE | Random 250 | Null |
| --- | --- | --- | --- |
| Train accuracy | $1 \pm 0$ | $1 \pm 0$ | $1 \pm 0$ |
| Test accuracy | $0.969 \pm 0.052$ | $0.781 \pm 0.117$ | $0.489 \pm 0.138$ |
| Sensitivity | $0.971 \pm 0.07$ | $0.779 \pm 0.175$ | $0.49 \pm 0.211$ |
| Specificity | $0.967 \pm 0.081$ | $0.786 \pm 0.173$ | $0.494 \pm 0.211$ |
| False positive | $0.029 \pm 0.087$ | $0.221 \pm 0.175$ | $0.51 \pm 0.211$ |
| False negative | $0.0033 \pm 0.081$ | $0.214 \pm 0.173$ | $0.506 \pm 0.211$ |

HD+ CAU and HD+ BA9 samples were applied to the random forests trained above. First, random forests built with the top 250 DE genes from (1) ranked by significance were used to predict the HD+ samples as either HD or C. We then built random forests with 250 randomly selected genes from (1), irrespective of significance, and performed classification of the HD+ samples. Finally, we built random forests with permuted sample labels and randomly selected genes to assess the basal predictive power under a null dataset. The results of these randomized random forest classifiers is included in Table 6.

### 4.8 t-statistic analysis of DESeq2 log fold changes

To identify genes that show different response between HD+ CAU and HD+ BA9 (3 vs 5) and early response genes from HD+ BA9 vs HD BA9 (1 vs 2), we developed a *t*-statistic methodology to quantify the difference between DESeq2 log2 fold change estimates while taking the uncertainty those estimates into account. DESeq2 implements a negative binomial generalized linear model, whose estimated coefficients are normally distributed. DESeq2 also reports the standard error of its log2 fold change estimates, enabling the calculation of a *t*-statistic corresponding to the confidence-adjusted difference in log2 fold change. Specifically, we calculate a *t*-statistic assuming both unequal sample sizes and unequal variance:

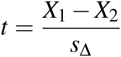

Where

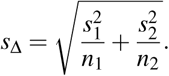

Here, *X*_1_ and *X*_2_ are the log2 fold change estimates from each comparison (e.g. (3) vs (5)), and *s*_1_ and *s*_2_ are the corresponding standard error estimates as reported by DESeq2. *n*_1_ and *n*_2_ are the number of samples total used for each analysis (e.g. for (3) vs (5), *n*_1_= 2 + 3 = 5 and *n*_2_ = 90 + 102 = 192, see Table 2). When assessing significance, the degrees of freedom is calculated using the Welch-Satterthwaite equation:

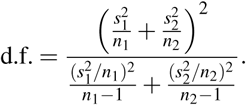

### 4.9 Comparison of DE Gene Lists and Enriched Gene Sets

*t*-statistics were calculated as described above for (3) vs (5) and (1) vs (2), where positive *t* corresponded to an increased log2 fold change in HD+ CAU over HD+ BA9 and HD+ BA9 over HD BA9, respectively. For (3) vs (5), genes were then ranked by descending *t*-statistic and analyzed for gene set enrichment with fgsea^6^. These GSEA results were then combined with those calculated for (1), (2), and (4). Significantly enriched gene sets at FDR < 0.05 were manually curated into 10 high level functional categories: Angiogenesis/Blood Brain Barrier (BBB), Apoptosis, Cell Cycle/Development, Cytoskeleton/Extracellular Matrix (ECM), Immune Response/Cancer, Metabolism, Neuron System, Protein Folding/Other, Signaling, and Transcription/Translation. Each gene set was also categorized into so-called Agreement Classes, an ordinal scale representing how specific the gene set is to HD+ CAU, as described in the results section. Combination of GSEA results, curation of gene sets, calculation of agreement classes, and plots from Figure 6 were made using python, jupyter lab, pandas, and matplotlib python libraries.

Gene sets within each functional category were also cast as a graph, where each node is a gene set, and edges between nodes were drawn if the gene sets shared more than 25% of their leading edge genes. Graph analysis was performed using python, networkx, and matplotlib. All analysis and figure code for this project are available at bitbucket.org/bubioinformaticshub/asymptomatic_hd_mrnaseq.

## Supporting information

Supplemental Table 1 - Sample Info

Supplemental Table 3 - Full fgsea results

Supplemental Table 4 - DE t-test results

Supplemental Table 2 - Full DESeq2 Results

## 5 Acknowledgements

We would like to thank the Jerry McDonald Huntington Disease Research Fund for supporting this work. The Genotype-Tissue Expression (GTEx) Project was supported by the Common Fund of the Office of the Director of the National Institutes of Health, and by NCI, NHGRI, NHLBI, NIDA, NIMH, and NINDS. The data used for the analyses described in this manuscript were obtained from dbGaP accession number phs000424.v7.p2 on 10/11/2017. Brain illustrations created by Freepik.

